# Homoploid hybridization signals due to ancestral subdivision: a case study on the D lineage in wheat

**DOI:** 10.1101/628685

**Authors:** Yunfeng Jiang, Zhongwei Yuan, Haiyan Hu, Xueling Ye, Zhi Zheng, Yuming Wei, You-Liang Zheng, You-Gan Wang, Chunji Liu

**Affiliations:** Triticeae Research Institute, Sichuan Agricultural University, Wenjiang, Chengdu 611130, China; CSIRO Agriculture and Food, St Lucia, Queensland 4067, Australia; College of Life Science and Technology, Henan Institute of Science and Technology, Xinxiang, Henan 453003, China; Science and Engineering Facility, Queensland University of Technology, Brisbane, QLD 4000

**Author notes:** These authors contributed equally to this publication (Yunfeng Jiang, Zhongwei Yuan).

**Keywords:** ancestral subdivision, ancestral variations, homoploid hybrid speciation, *Triticum-Aegilops*

## Abstract

Homoploid hybrid speciation has been reported in a wide range of species since the exploitation of genome sequences in evolutionary studies. However, the interference of ancestral subdivision has not been adequately considered in many such investigations. Using the D lineage in wheat as an example, we showed clearly that ancestral subdivision has led to false detection of homoploid hybridization signals. We develop a novel statistical framework by examining the changes in shared ancestral variations and infer on the likelihood of speciation due to genuine homoploid hybridization or ancestral subdivisions. Applying this to wheat data, we found that homoploid hybridization was not involved in the origin of the D lineage contrary to the now widely held belief. This example indicates that the significance of homoploid hybrid speciation is likely exaggerated. The underlying methodology developed in this study should be valuable for clarifying whether homoploid hybridization has contributed to the speciation of many other species.

## Introduction

Homoploid hybrid speciation (HHS) is the process of forming a new species between two donor species without a change in chromosome numbers. Such events have historically been considered rare, but this view has dramatically changed in recent years. Since the exploitation of sequences in evolutionary investigations, the involvement of homoploid hybridization (HH) has been proposed for the evolution and speciation of a wide range of species, including both plants and animals (Mallet 2007; Abbott et al. 2013; Sousa and Hey 2013; Payseur and Rieseberg 2016; Taylor and Larson 2019). However, the majority of the reported cases of HHS have been deduced from the analysis of sequence data only, while crucial evidence in support of such claims remains missing (Schumer et al. 2014). Accounts on the origins of the various progenitors of bread wheat (*Triticum aestivum* L.) serve as a typical example.

Bread wheat is an allohexaploid with three subgenomes (2n=6x=42; genome AABBDD). The three subgenomes are derived from three different diploid lineages (2x=2n =14): *Triticum urartu* (AA) (Dvorak et al. 1993), a close relative of *Aegilops speltoides* (BB) (Dvorak and Zhang 1990), and *Aegilops tauschii* (DD)(McFadden and Sears 1946). The phylogenetic histories of these diploid lineages have become the subject of controversy in recent years. Based on an analysis of genomic sequences, Marcussen et al. (2014) and Sandve et al. (2015) proposed the tantalising scenario that the ancestral D lineage originated from HH between ancient A and B lineages. By re-analysing data used by Marcussen et al. (2014) and chloroplast sequences from eight diploid and four polyploid wheat species in the *Triticum-Aegilops* complex, Li et al. (2015a, 2015b) argued for a more complex hybrid origin of the D lineage. Based on analyses of the evolutionary dynamics of transposon elements and mutations, El Baidouri et al. (2017) also reached the conclusion that the D lineage was derived from a complex history of multiple rounds of HH between ancient A and S (progenitor of the B subgenome) lineages as well as other relatives. Based on the analysis of transcriptome data, Glemin et al. (2018) concluded that pervasive HH events were involved in the evolution of these diploid wheat lineages, and that most of those belonging to the D lineage were derived from HHS between the A and B lineages.

To accommodate the discordant results obtained from the different studies, models for the evolution of the D lineage have become increasingly complicated. However, none of these models are strongly supported by biological evidence. First, the prohibitive genetic distances among the three diploid lineages of wheat known today make the prospects of producing any fertile diploid progeny between them unlikely. Even with modern techniques, it has not been possible to generate any viable diploid between the extant diploid A and D lineages, let alone between the more distant A and B lineages. Second, different chromosomal structures between these diploid lineages would have prevented the generation of fertile diploid hybrids between them. A large reciprocal translocation between chromosome arms 4AL and 5AL (Ma et al. 2013) exists in all extant species of the A lineage, including both *T. urartu* (King et al. 1994) and *T. monococcum* (Devos et al. 1995), but not in any species belonging to either the diploid B or D lineage. This translocation, if present prior to the assumed hybridization event(s) leading to the formation of the diploid D lineage, would make the production of any fertile diploid between the A and B lineages impossible. Third, restricted chromosomal recombination would be expected in hybrids involving distantly related genotypes (Ungerer et al. 1998; Buerkle et al. 2007) as well as genotypes with different chromosomal structures (Barb et al. 2014). However, large parental blocks from either the A or B genome were not detected on any of the D chromosomes (Marcussen et al. 2014; Glemin et al. 2018). Restricted recombination does not even exist in either of the chromosomal segments corresponding to the relative 4/5 translocation (supplementary fig.1).

The key evidence used in arguing for HHS of the D lineage is that, based on shared variations (gene trees, gene content, transposon elements (TE) and single nucleotide polymorphism (SNP)), the A and B lineages are more closely related to D individually than to each other (Marcussen et al. 2014; El Baidouri et al. 2017; Glemin et al. 2018). It is believed that, under the condition of random mating, shared variations concordant with the species tree should have the highest probability, while the other two discordant with the species tree should be equal in a bifurcating speciation. Any discrepancy with the above was accounted for by the involvement of HH (Hudson 1983; Tajima 1983). It is well known, however, that population subdivision occurs due to barriers such as geography or ecology (Slatkin and Pollack 2008), and that ancestral subdivision could lead to asymmetry in gene trees, resulting in the detection of false HH signals among descendant species based on many existing methods (Eriksson and Manica 2012; Eriksson and Manica 2014). Thus, the available results cannot rule out the possibility that the detected HH signals for the diploid wheat lineages could in fact be due to ancestral subdivision. The issue of how to distinguish between genuine HH and ancestral subdivision has been hotly debated in recent years (Durand et al. 2011; Eriksson and Manica 2012; Sankararaman et al. 2012; Yang et al. 2012; Eriksson and Manica 2014; Theunert and Slatkin 2017), and it remains very challenging. By developing and applying a new approach for detecting genuine HH, we re-evaluated whether the A and B lineages were indeed involved in the origin of the D lineage.

## Results and discussion

Three possible types of shared variations can be obtained from three different taxa (fig. 1A). Based on the times at which they occurred, shared variations can be placed into two classes. The first class are ancestral variations (AVs) which occurred before the differentiation of the earliest taxa among those under investigation. This class of variations can be grouped into three possible types under incomplete lineage sorting (ILS) (fig. 1A). Ancestral subdivision can lead to the unexpected distribution of AVs. The second class are variations which occurred between the differentiation of the first and last taxa under investigation. These variations reflect genuine evolutionary relationships, and they are thus termed phylogenetically informative variations (PIVs). The distribution patterns of PIVs should vary depending on whether HH was involved in the speciation of a given taxon (fig. 1A). Using the three diploid lineages of bread wheat as an example: If HH between the A and B lineages was not involved in the origin of the D lineage (the B(A,D) model), then PIVs should only be found in the shared variations between A and D (AD type). When a single event of HH was involved (the A(D)B model), PIVs should be present in both the AD and BD types (fig. 1A). Thus, differences in the patterns of PIV distribution can be reliably used to detect HH signals irrespective of whether ancestral subdivision was involved in the evolution of a given taxon.

**Figure 1.**
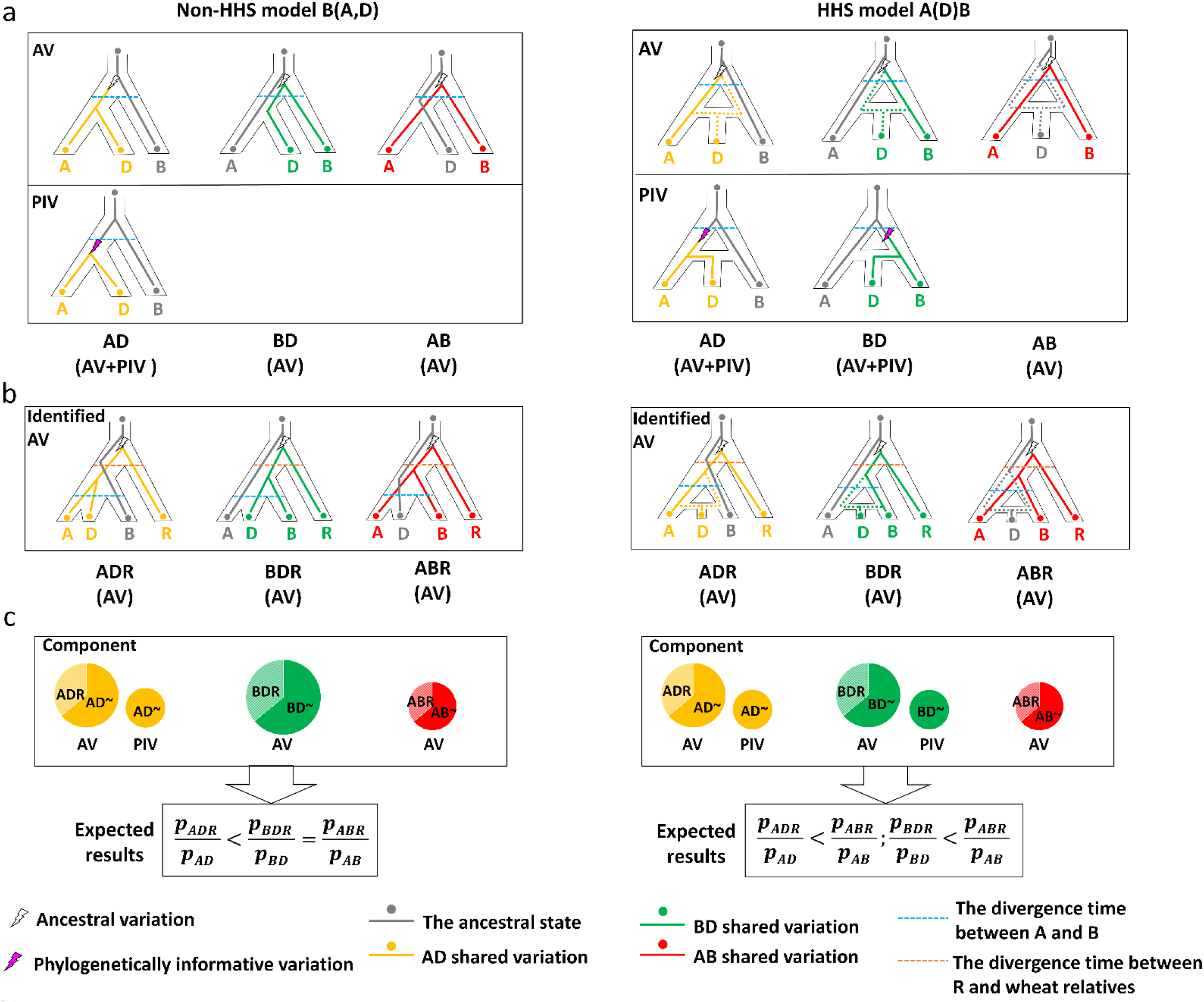
Difference in the inheritance patterns of shared variations between the HHS and non-HHS models for the speciation of the diploid D lineage in wheat. (A) Possible paths for the inheritance of ancestral variations (AVs) and phylogenetically informative variations (PIVs) for the models of non-HHS (left) or a single HH event (right). Dotted lines (grey, yellow and green) indicate possible paths of inherited variations. (B) AVs identified from the shared paths using rye (R) as the outgroup. (C) Difference in the expected results between the B(A,D) and A(D)B models. The probability (*p*) can be estimated from observed results from the shared variations.

Clearly, knowing the time at which a given variation occurred is the key to accurately identifying PIVs, but determining such times can be difficult. However, it should be possible to identify a proportion of AVs by introducing a suitable species as an outgroup. If the outgroup was differentiated earlier than any of the taxa under study, then shared variations among them should already exist in their common ancestor (fig. 1B). In other words, such variations should all belong to AVs. If the outgroup has a similar genomic relatedness to each of the taxa under investigation, then it would be possible to estimate the distribution of AVs. Thus, it should be feasible to deduce the PIV components based on AVs and use the information to detect genuine HH signals (fig. 1C and fig. 2A).

**Figure 2.**
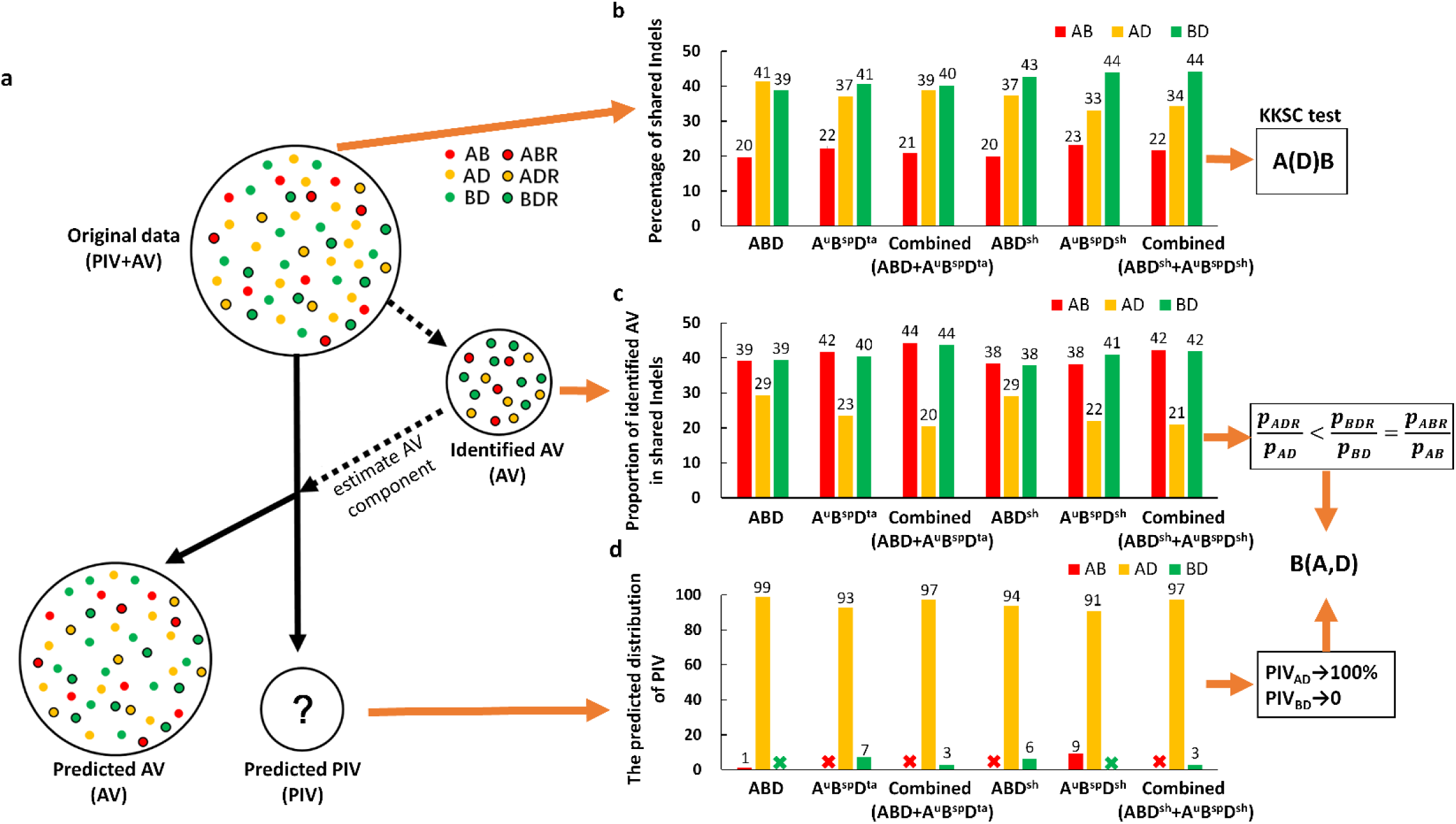
Phylogenic relationships among the diploid lineages of A, B and D in wheat inferred from shared variations. Analyses were conducted on six data sets from the different combinations of the three lineages: A lineage including A subgenome (A) and *T. uratu* (A^u^); B lineage including B subgenome (B) and *Ae. speltoides* (B^sp^); D lineage including D subgenome (D), *Ae. tauschii* (D^ta^) and *Ae. Sharonensis* (D^sh^). (A) An illustration of the approach used to indirectly detect PIVs from the original data. (B) Percentages of shared variations (Indels) in AB (red bars), AD (yellow bars) and BD (green bars) based on the original data. KKSC insertion significance test (Kuritzin et al. 2016) support the A(D)B model (supplementary table 3). (C) The proportion of AVs identified from the shared Indels. The values of AB (red bars), AD (yellow bars) and BD (green bars) were calculated based on the numbers of shared Indels in ABR/AB, ADR/AD and BDR/BD, respectively. The Chi-squared test against these data supported 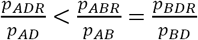 (supplementary table 4). (D) Percentages of shared variation between AB, AD and BD after removing the estimated components of AVs. The majority of AVs could be removed from the original data based on their distributions until one of the three shared types reached zero (the symbol of“×”).

We assessed rye (R genome) as an outgroup to identify AVs for the three diploid wheat lineages. Rye diverged earlier (about 4.5 MY) than the diploid wheat lineages (Huang et al. 2002; Gornicki et al. 2014), and the genomic relatedness between it and the three wheat lineages was similar (supplementary fig. 2 and supplementary table 1). These characteristics indicate that the distribution of AVs among the three diploid wheat lineages can be estimated based on their presence or absence in rye (supplementary fig. 3).

We thus conducted an analysis based on shared Indels in 3,771 orthologous genes identified from the three diploid wheat lineages: A lineage, including the A subgenome of bread wheat (genome A) and *T. uratu* (genome A^u^); B lineage, including the B subgenome (B) and *Ae. speltoides* (B^sp^); and D lineage, including the D subgenome(D), *Ae. tauschii* (D^ta^) and *Ae. sharonensis* (D^sh^). Different combinations of the three lineages yield a total of six different datasets, including ABD, A^u^B^sp^D^ta^, combined(ABD+A^u^B^sp^D^ta^), ABD^sh^, A^u^B^sp^D^sh^ and combined(ABD^sh^+A^u^B^sp^D^sh^) (supplementary table 2). Initially, shared Indels in the different datasets were identified using barley to infer their ancestral states. This analysis detected 881, 711, 592, 730, 645 and 514 shared Indels, respectively, from each of the datasets. Similar to data reported earlier (Marcussen et al. 2014; El Baidouri et al. 2017), these original data suggest that the A and B lineages are more closely related individually to D than to each other, and that BD is closer than AD in most of the combinations (except for ABD) (fig. 2b and supplementary table 2). A KKSC insertion significance test (Kuritzin et al. 2016) against these data strongly (*p* < 0.01) suggested HHS of the D lineage from A and B based on each of the datasets (fig. 2b and supplementary table 3). However, as mentioned earlier, using such data cannot eliminate the interference of ancestral subdivisions.

We then used the R genome to identify AVs from each of these datasets. This analysis found that the proportions of ABR in AB and BDR in BD are very similar. However, the proportion of ADR in AD is significantly lower than the other two types (fig. 2C and supplementary table 4). Clearly, the proportion of AVs from shared variations containing AVs should only be higher than those containing both AVs and PIVs (fig. 1C). Thus, strong signals of PIVs were present in the AD type but not in the BD type, which was expected from a non-HH model of B(A,D). To determine the distribution of PIVs, we simulated the effects of removing AVs from the original data based on the observed ratios of these variations until one of the three shared types reached zero (fig. 2A). If the observed ratios are similar to the theoretical distribution of the AVs, then the removed part should predominantly originate from AVs, while the remainder should mainly contain PIVs. This method should thus provide a good estimate of the distribution of PIVs. Here, all datasets showed that the percentages of the AD type were very high (91%∼99%) in the predicted distribution of PIVs (fig. 2D). However, few of the shared variations in the BD type were retained in most of the datasets (fig. 2D), indicating that they predominantly originated from AVs. Again, these results all support the non-HHS model of B(A,D) and are not what should be expected from the HHS model of A(D)B.

We then further estimated the optimal parameters for both the HHS and non-HHS models using the shared variations by maximising the multinomial log-likelihood. We tested which of the models could be accepted based on the discrepancy between the observed and expected data for each of them. The results showed that significant differences were not detected from the datasets for any of the combinations under the non-HHS model B(A,D) (fig. 3A and table 1). However, the HHS model A(D)B (with equal contributions of A and B) (Marcussen et al. 2014) was strongly rejected (fig. 3b and supplementary table 5). Further, the parameters from the non-HHS model B(A,D) provided a clear picture on how AVs led to the detection of false HH signals. High proportions of AVs (varying from 79% to 91%) were present in the original data, and their distributions did not meet the ratios of AD>AB=BD, as should be expected from ILS under random mating. Understandably, the interactions between the high proportion and unexpected distribution of AVs could easily obscure the effect of PIVs in the original data, resulting in the A and B lineages appearing to be more closely related to the D lineage individually than to each other.

**Figure 3.**
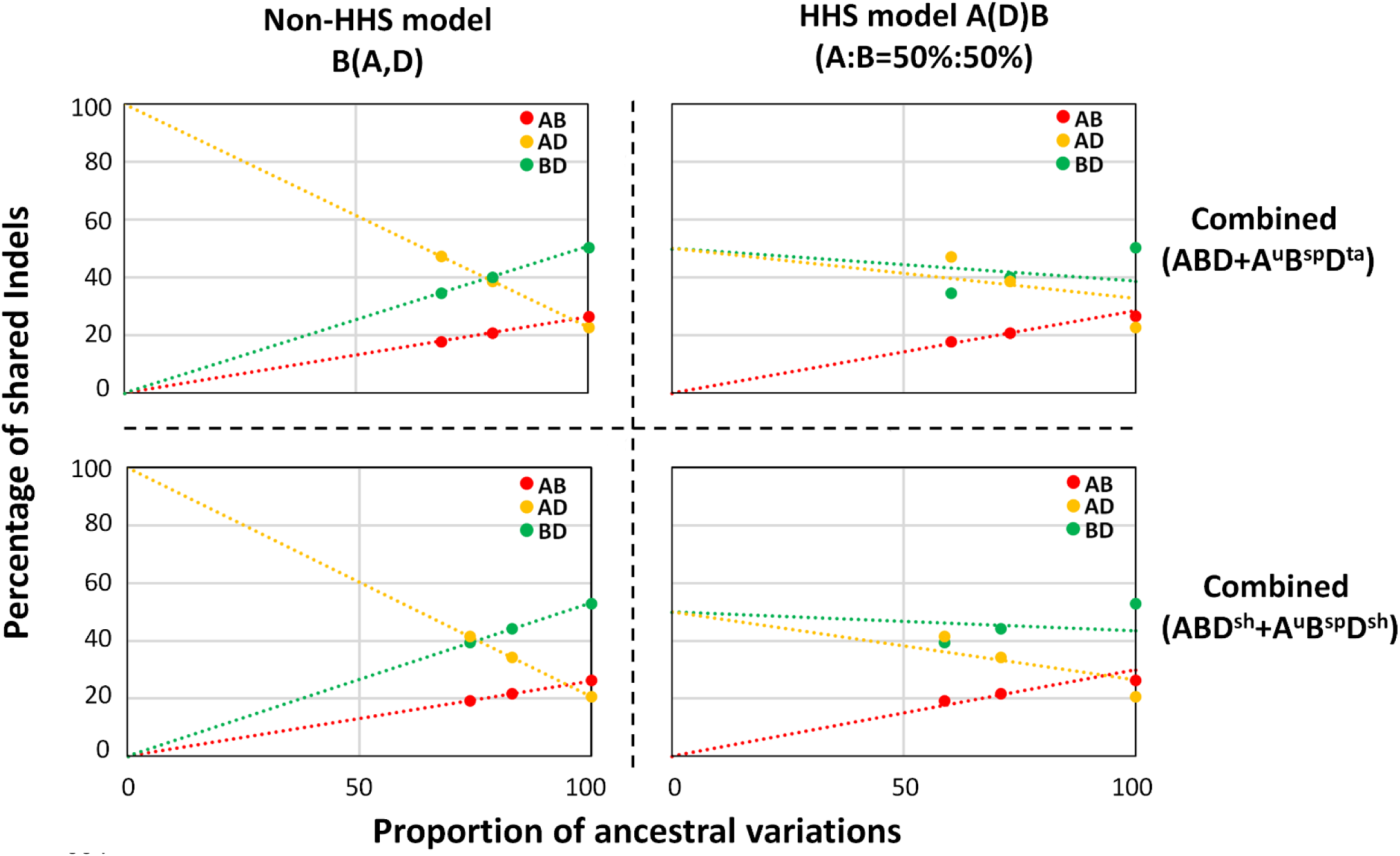
Fitness comparison between the HHS and non-HHS models based on expected and detected ratios of shared variations. Two combined data sets were used to assess the HHS model A(D)B (with equal contributions) (Marcussen et al. 2014) and the non-HHS model B(A,D). The dots show the detected results from the shared variations (supplementary table 2), and the dotted lines the expected results based on either of the models.

**Table 1.** The statistical test for the non-HHS model B(A,D)

| Data set<sup>a</sup> | Proportion of<br>AVs in<br>shared Indels | Predicted distribution of AVs |  |  | Proportion of PIVs<br>(only AD) in shared<br>Indels | Significant test<br>(p-value)<sup>b</sup> |
| --- | --- | --- | --- | --- | --- | --- |
|  |  | AB | AD | BD |  |  |
| ABD | 89.6% | 22.0% | 34.5% | 43.4% | 10.4% | 0.951 |
| A<sup>u</sup>B<sup>sp</sup>D<sup>ta</sup> | 84.2% | 26.4% | 25.3% | 48.3% | 15.8% | 0.821 |
| Combined<br>(ABD+A<sup>u</sup>B<sup>sp</sup>D<sup>ta</sup>) | 79.2% | 26.4% | 22.8% | 50.8% | 20.8% | 0.905 |
| ABD<sup>sh</sup> | 91.2% | 21.9% | 31.2% | 46.9% | 8.8% | 0.912 |
| A<sup>u</sup>B<sup>sp</sup>D<sup>sh</sup> | 85.2% | 27.1% | 21.4% | 51.5% | 14.8% | 0.581 |
| Combined<br>(ABD<sup>sh</sup>+A<sup>u</sup>B<sup>sp</sup>D<sup>sh</sup>) | 82.9% | 26.1% | 20.7% | 53.3% | 17.1% | 0.931 |

<sup>a</sup> A lineage including A subgenome (A) and *T. uratu* (A<sup>u</sup>); B lineage including B subgenome (B) and *A. speltoides* (B<sup>sp</sup>); D lineage including D subgenome(D), *A. tauschii* (D<sup>ta</sup>) and *A.* *sharonensis*(D<sup>sh</sup>).
<sup>b</sup> The null hypothesis is based on the non-HHS model B(A,D). That the p-values for none of the combinations are significant indicates that there is no evidence against the model.

In addition to shared variations, estimated divergence times (EDTs) based on nuclear genome sequences that contradict a treelike phylogeny for the three diploid lineages of bread wheat (fig. 4A) were also used as evidence to argue for the HHS of the D lineage (Marcussen et al. 2014). However, it is known that AVs could lead to an overestimation of divergence times (Charlesworth 2010). EDTs for the three diploid lineages of bread wheat based on the nuclear genomes (Marcussen et al. 2014) are more than double those based on the chloroplast genomes (Gornicki et al. 2014) (fig. 4A). The differences in the levels of overestimation among the three lineages based on the nuclear sequences can be explained by the different levels of AVs among them (fig. 4B). As the proportion of AVs in the BD type is much larger than that in the AB type (table 1), it is expected that the coalescent EDT(A-B) should be much higher than the coalescent EDT(B-D) based on the nuclear sequences (fig. 4B). Thus, the EDTs of the three lineages based on the nuclear genomes cannot be used as evidence to argue for the HHS of the D lineage. As chloroplast genomes are predominantly maternally inherited, variations in them should not be significantly affected by AVs. As a result, EDTs from the chloroplast sequences should be more reliable. Based on these understandings and the available results, we revised the model for the evolution of the three diploid lineages of bread wheat (fig. 5).

**Figure 4.**
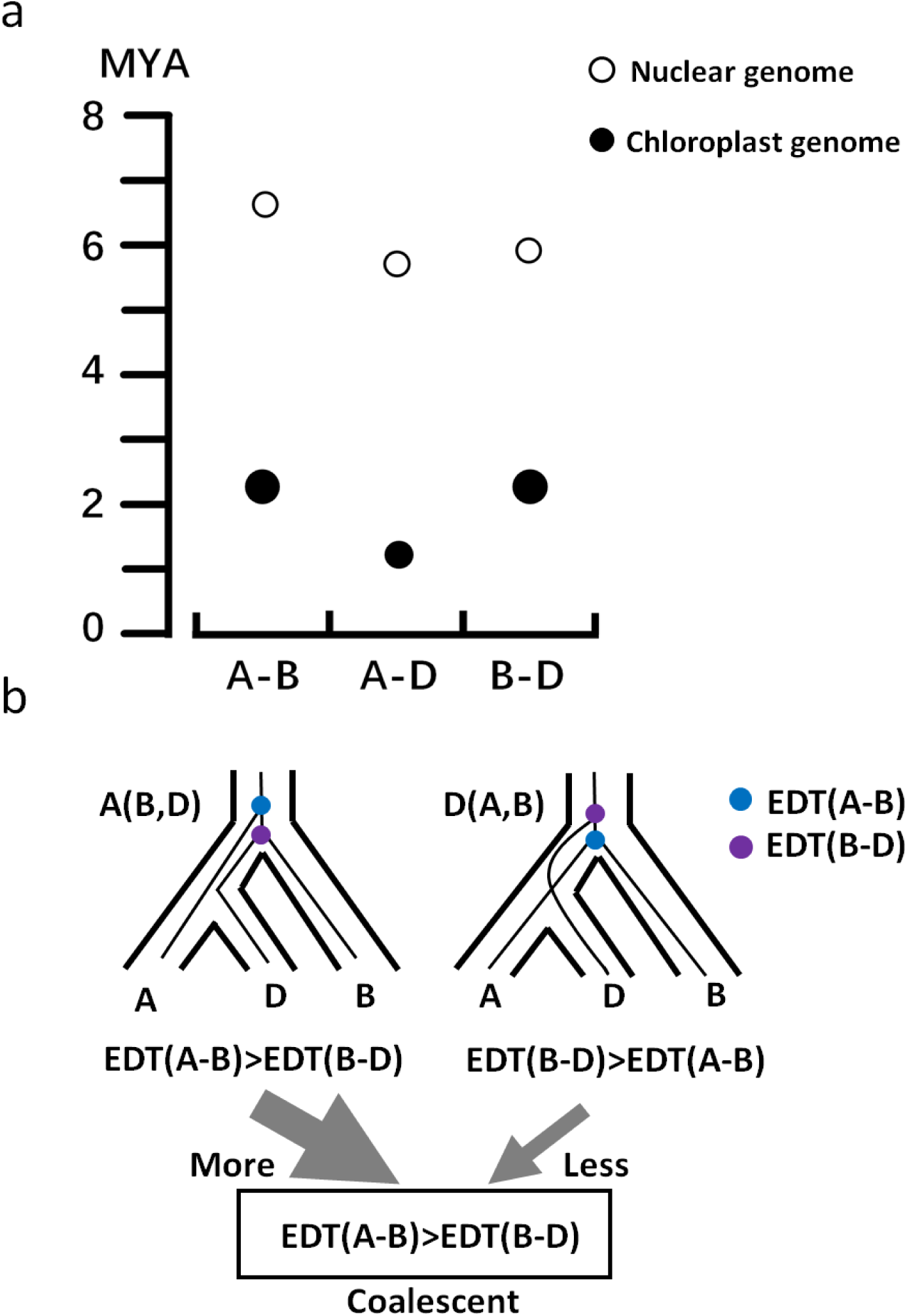
Estimated divergence times (EDTs) under the influence of ancestral variations (AVs). (A) Difference in EDTs between the uses of nuclear (Marcussen et al. 2014) and chloroplast (Gornicki et al. 2014) sequences. Diameters of the dots indicate the confidence interval. (B) Overestimation of divergence times due to AVs. The gene trees of D(A,B) and A(B,D) originated from AVs likely resulted in overestimations of the coalescent EDT of the three lineages. Since the proportion of A(B,D) is much larger than that of D(A,B), the coalescent EDT(A-B) would likely be more significantly overestimated in comparison to the coalescent EDT(B-D).

**Figure 5.**
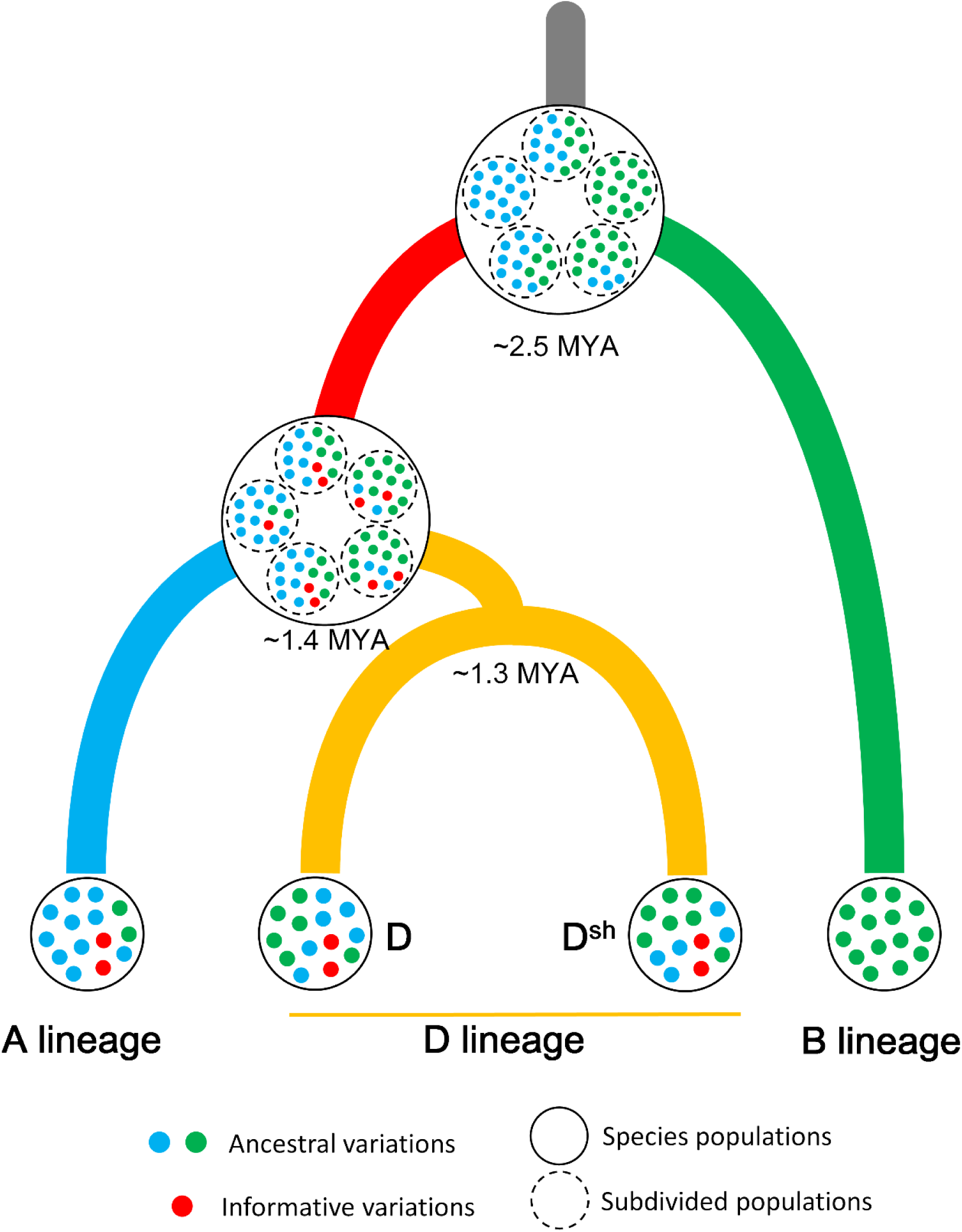
Evolutionary relationships among the three diploid lineages of bread wheat. The divergence times were inferred from chloroplast sequences (Gornicki et al. 2014).

Using the three diploid lineages of bread wheat as an example, we demonstrated that the interference of ancestral subdivision could pose a huge problem in detecting genuine HH signals in speciation. The diploid D lineage of bread wheat is only one of the many taxa for which the possibility of HHS has been deduced from sequence data (Schumer et al. 2014). Although the possibility that HHS has played critical roles in the evolution of some species may exist, such possibility is now clearly excluded in the study of D lineage of wheat, we therefore believe that its importance has likely been exaggerated. The new approach discussed in this publication not only provides more meaningful results for the evolution of the D lineage of bread wheat but should also be invaluable in re-assessing other species for which HHS has been deduced based on sequence data.

## Materials and Methods

### Genome sequences used in the study

Multiple sets of genome sequences were used in this study. Genome sequences and gene models of bread wheat based on IWGSC RefSeq assembly v1.0 (IWGSC 2014) were downloaded from http://www.wheatgenome.org/. Genome sequences of *Triticum urartu* (AA) (Ling et al. 2013) and *Aegilops tauschii* (DD) (Jia et al. 2013) were downloaded from http://plants.ensembl.org/. The diploid species of B lineage (*Ae. speltoides* (SS)) and D lineage (*Ae. sharonensis*) were downloaded from http://www.wheatgenome.org/. The genome sequences and gene models of barley (*Hordeum vulgare* L; 2n=2x=14; genome HH) (Mascher et al. 2017) were obtained from http://plants.ensembl.org/. Rye (*Secale cereal*, genome RR) was used to identify ancestral variations in the diploid progenitors of bread wheat, and its genome sequences (Bauer et al. 2017) were downloaded from http://webblast.ipk-gatersleben.de/ryeselect/.

### Genomic compositions of the D lineage of bread wheat based on orthologous gene trees using the maximum likelihood method

To investigate whether it originated from HHS (Marcussen et al. 2014) between the A and B lineages, genomic compositions of the D lineage from each of the putative parents were measured based on orthologous gene trees using the maximum likelihood method (Stamatakis 2014). Putative orthologous genes were identified from predicted proteins in barley and the three subgenomes using the PorthoMCL (Tabari and Su 2017) software with the default parameters.

The predicted coding sequences (CDS) from the putative orthologous genes in barley were then compared with those in the three subgenomes of bread wheat using BLASTn. Orthologous clusters were screened with a similarity > 85% for at least 50% of the lengths among them. Apart from the regions involved in the reciprocal translocation between chromosome arms 4AL and 5AL, only clusters containing strictly four genes belonging to a single homoeologous group were considered as robust orthologous quadruplets (supplementary data 1). Chromosomal locations of 4A/5B/5D/5H and 5A/4B/4D/4H were used to identify the regions corresponding to the translocation (supplementary data 2). CDS from the identified orthologous quadruplets were aligned using MAFFT (Katoh and Standley 2013) with default parameters. They were used to reconstruct individual-gene maximum likelihood phylogenies using barley as the outgroup, with RAxML (Stanmatakis 2014) based on the GTRGAMMA substitution model. Bootstrap support was calculated from 200 replicates. Phylogeny trees with bootstrap support of less than 50% were removed, and the remainder were further grouped to identify their origins. The CDS were then ordered according to their physical locations on the D subgenome of bread wheat. To measure approximate sizes of the parental blocks (Ungerer et al. 1998), physical sizes of consecutive genes of the same parentage were calculated (the distance spanned by all consecutive genes from the same parental species plus one-half of the distance to the nearest genes from the other species). Blocks at the ends of each group were not included because of the difficulty involved in determining their sizes.

### Genomic relationships between rye and the diploid progenitors of bread wheat

Compared with barley, rye could be considered as a closer outgroup to determine the evolutionary relationships among the three diploid donors of bread wheat (Glemin et al. 2018). To assess this feasibility, we re-examined their relationships based on genome sequences. First, the genomic relatedness among the three diploid progenitors of bread wheat and rye was visualised with the rooted phylogenetic network based on the sequence data used by Marcussen et al. (2014). These data included the three subgenomes of bread wheat and the genomes of *T. uratu, T. monococcum, Ae. speltoides, Ae. tauschii, Ae. sharonensis* and *H. vulgare*. They were analysed against the CDS database of rye (Bauer et al. 2017). This analysis obtained 192 consensus sequences from the rye genome (supplementary data 3). The orthologous sequences were used to reconstruct individual-gene maximum likelihood phylogenies with barley as the outgroup using the RAxML (Stamatakis 2014) with the GTRGAMMA substitution model. Bootstrap support was calculated from 200 replicates. A Neighbour-Net was constructed from the average pairwise distances among the 192 gene trees using SplitTree 4 (Huson 1998). Second, the genomic relatedness among the three diploid donors of bread wheat and rye were further assessed using shared single nucleotide polymorphisms (SNPs). This assessment was based on 1,614 orthologous genes identified from the homoeologous CDS among the genomes of rye, barley and the three subgenomes of bread wheat (supplementary data 4). The predicted CDS of the five orthologous quintuplets were aligned using MAFFT (Katoh et al. 2013) with default parameters. SNPs were automatically detected using SNP-sites (Page et al. 2016). Indels were eliminated to avoid differences caused by different transcripts. Four different datasets, including A/B/R, A/D/R, B/D/R and A/B/D, were analysed, and the ancestral states were inferred based on the barley genome sequences. Only variations that split the four taxa into two groups of two were considered so that three types of shared variations could be obtained. For example, the three types of shared variations for the A/B/R dataset are AB, AR and BR types.

### Identification of shared Indels

Indels have previously been widely utilised to assess evolutionary relationships. We focused only on those Indels from genes which could be found from each of the three subgenomes of bread wheat and from the genome of barley in this study. A total of 3,771 such genes (orthologous quadruplets) were found (supplementary data 1) and used for further analyses. Sequences of the orthologous genes for the three subgenomes of bread wheat (ABD) were extracted from the wheat genome based on their locations from the gene models (IWGSC 2014). Their orthologous sequences in *T. urartu* (A^u^), *Ae. speltoides* (B^sp^), *Ae. tauschii* (D^ta^) and *Ae. sharonensis* (D^sh^) were identified based on BLAST analysis against the sequences of barley. Initially, four different datasets from different combinations of these genomes were obtained. These included ABD, A^u^B^sp^D^ta^, ABD^sh^ and A^u^B^sp^D^sh^. These quadruplets of orthologous genes were aligned using MAFFT with default parameters. Putative Indels with lengths of ≥ 5 bp were screened using RELINDEL (Ashkenazy et al. 2014) with default parameters. Further, three types of the shared Indels were manually detected based on the following rules. Briefly, the presence or absence of an Indel in the outgroup (barley) was treated as the ancestral state (as 0), and the alternative was treated as the mutated state (as 1). Considering the order of H|A|B|D, we have 0|1|1|0 for AB shared Indels, 0|1|0|1 for AD shared Indels, and 0|0|1|1for BD shared Indels.

When multiple Indels were present in a single gene, those with the same sequence were recorded only once. To decrease sampling bias from a single dataset, two combined datasets were generated from the common variations shared by ABD+A^u^B^s^D^ta^ and ABD^sh^+A^u^B^s^D^sh^. Thus, subsequent analyses were performed using a total of six different datasets of shared variations (including ABD, A^u^B^s^D^ta^, ABD^sh^, A^u^B^s^D^sh^, combined ABD+A^u^B^s^D^ta^ and combined ABD^sh^+A^u^B^s^D^sh^). A KKSC insertion significance test (Kuritzin et al. 2016) was conducted against these variations.

The shared Indels were then put into two groups based on their presence or absence in the rye genome. For those present in R, we have 0|1|1|1|0 (in the order of H|R|A|B|D) for ABR shared Indels, 0|1|1|0|1 for ADR shared Indels, and 0|1|0|1|1 for BDR shared Indels. For those absent in R, we have 0|0|1|1|0 for AB∼ shared Indels, 0|0|1|0|1 for AD∼ shared Indels, and 0|0|0|1|1 for BD∼ shared Indels. These data are listed in supplementary data 5.

### Distinguishing homoploid hybridization and ancestral subdivision from shared variations

Shared variations could be divided into two classes based on the times at which they occurred: (1) Ancestral variations (AVs), which occurred before the differentiation of the earliest species under investigation; and (2) phylogenetically informative variations (PIVs), which occurred between the differentiation of the first and last species of concern. With three taxa, there are three possible patterns of shared variations from AVs under ILS. However, the patterns of PIV distributions should differ depending on whether HH was involved. Thus, PIVs can be unambiguously used to reconstruct evolutionary relationships. However, it is difficult to precisely identify PIVs from the mix due to the difficulty involved in determining the precise time at which a given variation occurred. Here, we propose an alternative approach which, by estimating the distribution of AVs, allows the indirect detection of PIVs.

### The distribution of AVs

Suppose the total number of shared variations consisting of two classes, AVs and PIVs, we have *p*_*AV*_ + *p*_*PIV*_ = 1. As AVs are distributed among AB, AD and BD, we have *p*_*AV*_ = *p*_*AV*(*AD*)_+*p*_*AV*(*BD*)_+*p*_*AV*(*AB*)_. With the use of a suitable outgroup, we can estimate the probability of AV distribution. The outgroup must satisfy the following three criteria: (1) it diverged earlier than any of the diploid lineages of bread wheat from a common ancestor but is close enough and shares a proportion of AVs with them; (2) gene flows between the outgroup and the taxa under investigation have not occurred. Thus, shared variations between it and the targeted taxa should be predominantly derived from AVs; and (3) the outgroup has similar genomic relatedness to the taxa under investigation. In other words, inherited AVs by the outgroup should not be biased towards any one of them. Rye seems to be a suitable candidate as the outgroup for the diploid lineages of bread wheat, as it satisfies these criteria (supplementary fig. 2 and supplementary table 1). Based on whether they are present in rye, we can obtain a dataset containing AVs only. This dataset includes all shared variations present in ADR, BDR and ABR. Understandably, due to incomplete lineage sorting (ILS), a proportion of the AVs occurring prior to the divergence of rye (AVs^before R^) would show ancestral states in the R genome. Thus, not all AVs^before R^ could be detected using rye as the outgroup. Supposing that *ω* is the proportion of AVs^before R^ in all AVs, then the probability of AD, BD and AB types derived from AVs^before R^ could be represented as *ωp*_*AV*(*AD*)_, *ωp*_*AV*(*BD*)_ and *ωp*_*AV*(*AB*)_, respectively. Supposing that *γ*_1_, *γ*_2_ and *γ*_3_ are the proportions of shared variations between rye and those in AD, BD or AB from AVs^before R^, respectively, then we have *pADR*=*γ*_1_*ωpAV*(*AD*), *pBDR*=*γ*_2_*ωpAV*(*BD*) and *pABR* =*γ*_3_*ωpAV*(*AB*). Here, *γ*_1_, *γ*_2_ and *γ*_3_are determined by the genomic relatedness between rye and the three diploid lineages of wheat. As discussed earlier, *γ*_1_, *γ*_2_ and *γ*_3_ should be very close to each other. We therefore assumed that *γ*_1_ = *γ*_2_ = *γ*_3_ = *γ*, and that the distribution of AVs could therefore be estimated from the identified AVs (i.e., *pADR*: *pBDR*: *pABR* = *pAV*(*AD*): *pAV*(*BD*): *pAV*(*AB*)).

### Method 1: Detecting PIV signals from shared variations

As the proportions of inherited variations from AV^before R^ between rye and those in AD, BD or AB types are similar, we have

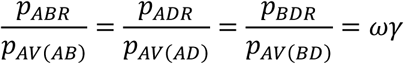

As all shared variations in AB should originate from AVs in our models, we have *p*_*PIV*(*AB*)_ =0 and *p*_*AB*_ = *p*_*AV(AB)*_. Thus, 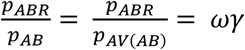 If AD or BD shared variations include PIVs (*p*_*PIV*(*AD*)_ > 0 or 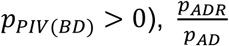 or 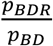 will be less than 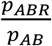 We thus tested the two hypotheses to detect PIV signals from the shared variations in AD and BD, respectively.

#### PIV signals in AD

The hypothesis test here is *H*_0_: *p*_*PIV*(*AD*)_ = 0 versus *H*_1_: *p*_*PIV*(*AD*)_ > 0.

The hypothesis *p*_*PIV*(*AD*)_ = 0 is equivalent to 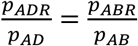, and columns (AD and AB) and rows (present in R and absent in R) are independent, as in the analysis of a contingency table. Rejection of *H*_*0*_ indicates that 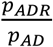 is less likely than 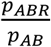, i.e., there is a significant number of PIV signals in AD.

#### PIV signals in BD

The hypothesis test is *H*_0_: *p*_*PIV*(*BD*)_ = 0 versus *H*_1_: *p*_*PIV*(*BD*)_ > 0. The hypothesis *p*_*PIV*(*BD*)_ = 0 is equivalent to 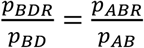. Rejection of *H*_*0*_ indicates 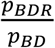 is less likely than 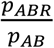, i.e., there is a significant number of PIV signals in BD.

Here, 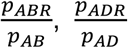 and 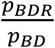 could be estimated from the observed values of 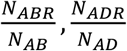 and 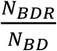, respectively. The differences among 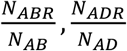 and 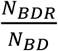 were compared using a four-fold table with chi-squared tests.

In summary, if significant PIV signals are detected in the AD type but not in the BD type, then the non-HHS model B(A,D) should be accepted. On the other hand, if significant PIV signals are detected in both the AD and BD types, then the HHS model A(D)B should be accepted.

### Method 2: Differentiating the HHS- and non-HHS models based on the discrepancy between observed and expected ratios of shared variations

Using the R genome as an outgroup, we can divide the original data into AVs and mixed variations which contain both AVs and PIVs. We can express their distributions (original data, AVs and mixed variations) in the AB, AD and BD types as *y*_*AB*_ = β_*AB*_*x* + α_*AB*_, *y*_*AD*_ = β_*AD*_ *x* + α_*AD*_ and *y*_*BD*_ = β_*BD*_ *x* + α_*BD*_, respectively, where α_*AB*_, α_*AD*_ and α_*BD*_ are the proportions of the PIVs among the shared variations in the AB, AD and BD types. We thus have *α*_*AB*_ + *α*_*AD*_ + *α*_*BD*_ = 1; *x* is the proportion of AVs in the shared variations. Note that βs are fixed rate parameters: A negative value suggests that the proportion of the shared variations (*y*) will increase gradually with decreasing *x* (when removing AVs based on the presence or absence of a given variation in the outgroup). Here, *y*_*AB*_, *y*_*AD*_ and *y*_*BD*_ vary as *x* changes, while keeping *y*_*AB*_ + *y*_*AD*_ + *y* _*BD*_ = 1. In the original data, we have 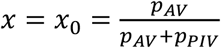, which is smaller than 1. In the mixed variations: Supposing *ω*γ is the proportion of removed AVs, the value of *x* becomes 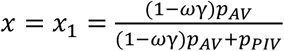, which is smaller than *x*_0_. In the AV data, all shared variations are AVs; thus, the corresponding *x* is 1.

Under the non-HHS model B(A,D), the expected *α*_*AD*_ should be close to 1. We therefore tested the hypothesis *H*_0_: *α*_*AD*_ = 1 versus *H*_1_: *α*_*AD*_ ≠ 1. Under the HHS model A(D)B, the expected *α*_*AD*_ ≃ *α*_*BD*_ ≃ 0.5, i.e., roughly equal contribution were expected from each of parental lineages to the D (Marcussen et al. 2010). We can therefore also test the hypothesis of *H*_0_: *α*_*AD*_ = *α*_*BD*_ = 50%.. The test statistic was constructed based on (Observed-Expected)^2/Expected as in the analysis of contingency table (Agresti 2018), and the expected values were obtained by maximising the multinomial log-likelihood and incorporating the constraints under the null hypothesis.

## Acknowledgements

The work was supported by the Commonwealth Scientific and Industrial Organization (CSIRO), Australia (Project code: R-10191-01). Y.J., Z.Y., X.Y. are grateful to the Sichuan Agricultural University and the China Scholarship Council for funding his visit to CSIRO. H.H. is grateful to Henan Institute of Science and Technology and CSC for supporting her visit to CSIRO. The authors are grateful to Dr. Donald Gardiner for his constructive suggestions during the preparation of the manuscript.

---

**Supplementary Figure 1.**
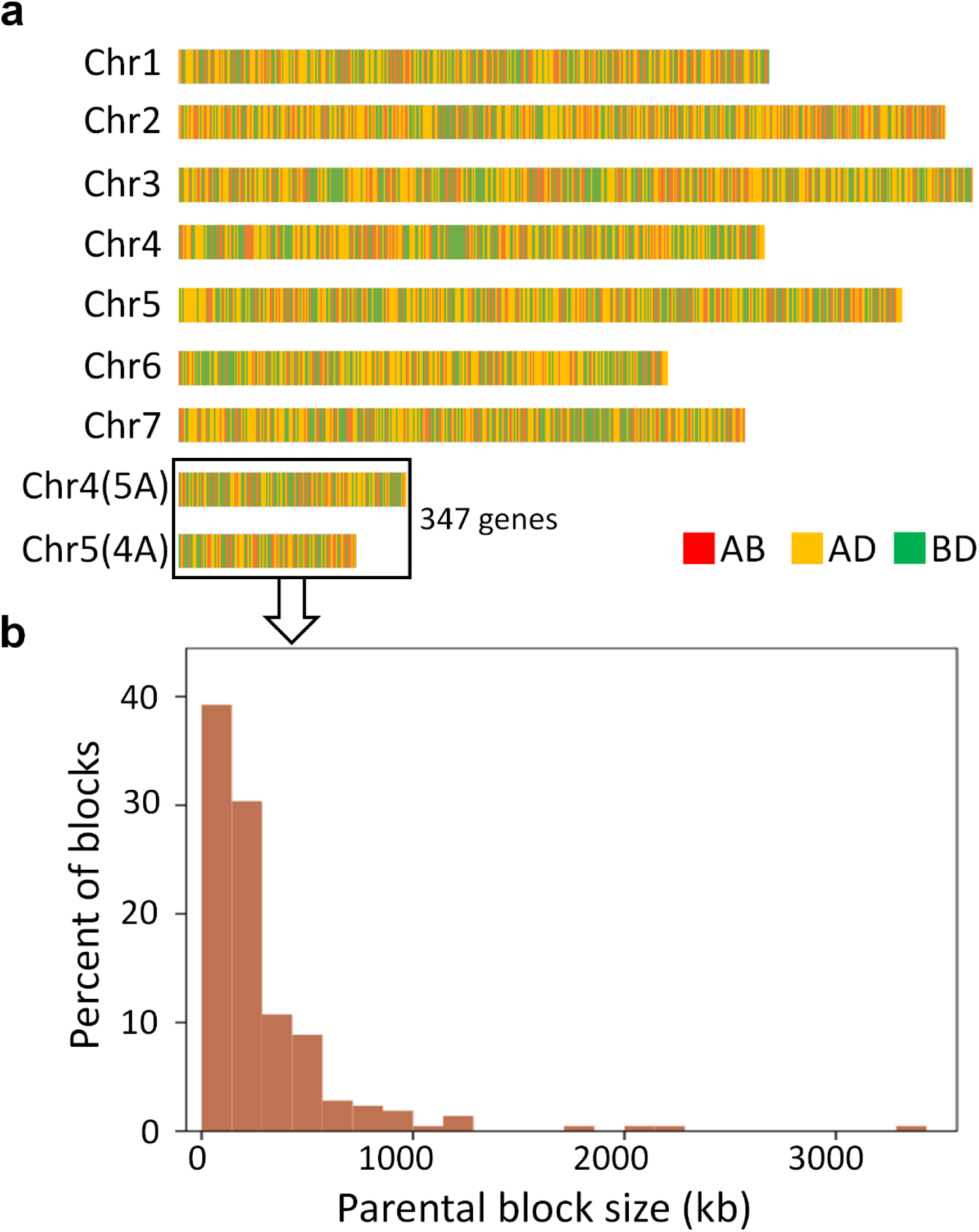
Distributions of parental gene blocks for the three subgenomes of bread wheat based on gene trees. (a) RAxML results based on the use of 3,771 genes on the seven chromosomes and 347 genes on the segments related to the 4AL/5AL translocation. AB, AD and BD represent the gene trees of D(A,B), B(A,D) and A(B,D), respectively. They were ordered according to the physical map of the D subgenome. (b) Size distribution of parental blocks in the two regions involved in the relative 4AL/5AL translocation.

**Supplementary Figure 2.**
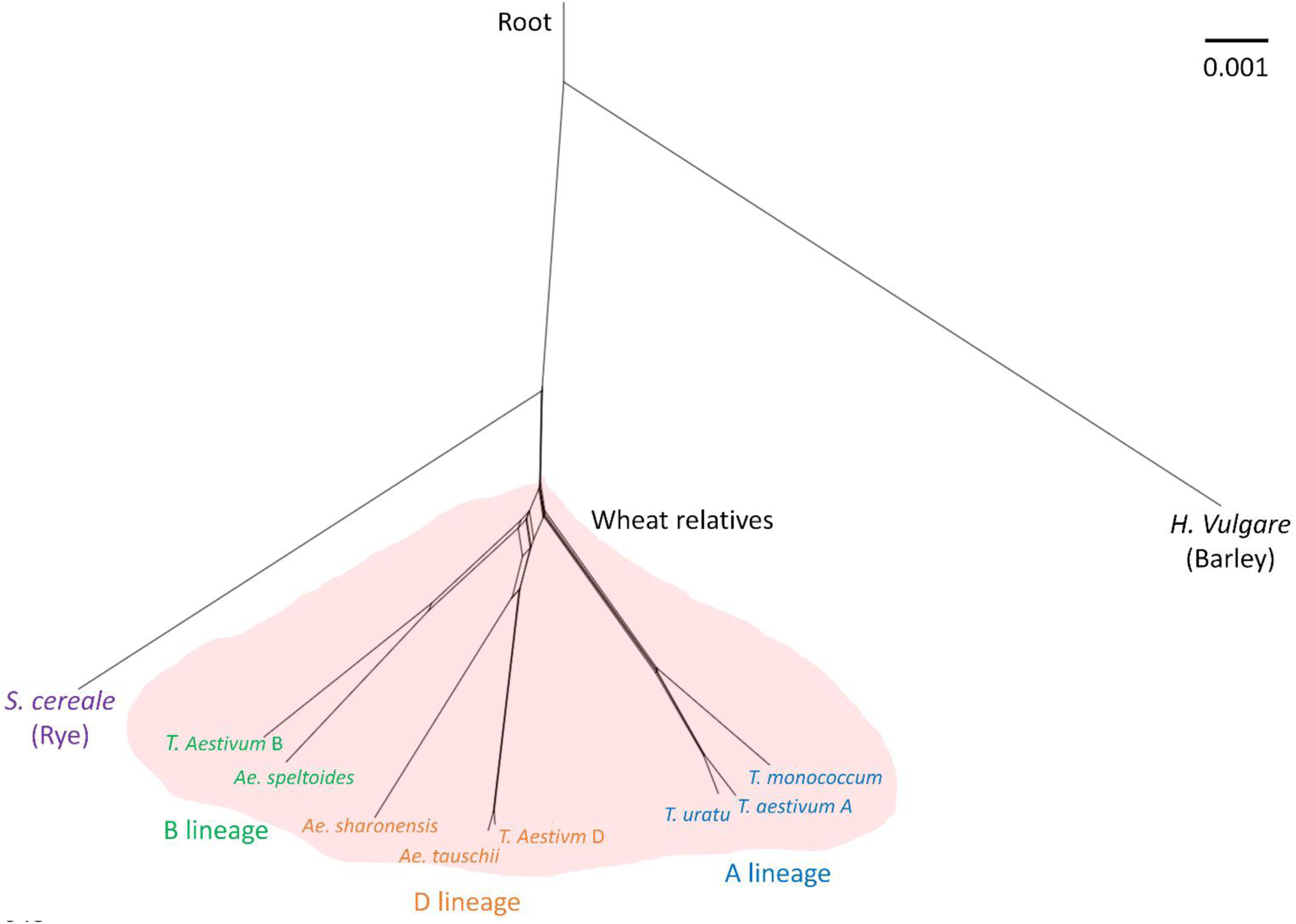
The rooted phylogenetic network constructed from the RAxML results based on the analysis of split networks using the 192 orthologous genes. It shows clearly that rye diverged earlier than the three diploid lineages of bread wheat and no gene flow occurred between them after the divergence of the A, B and D lineages.

**Supplementary Figure 3.**
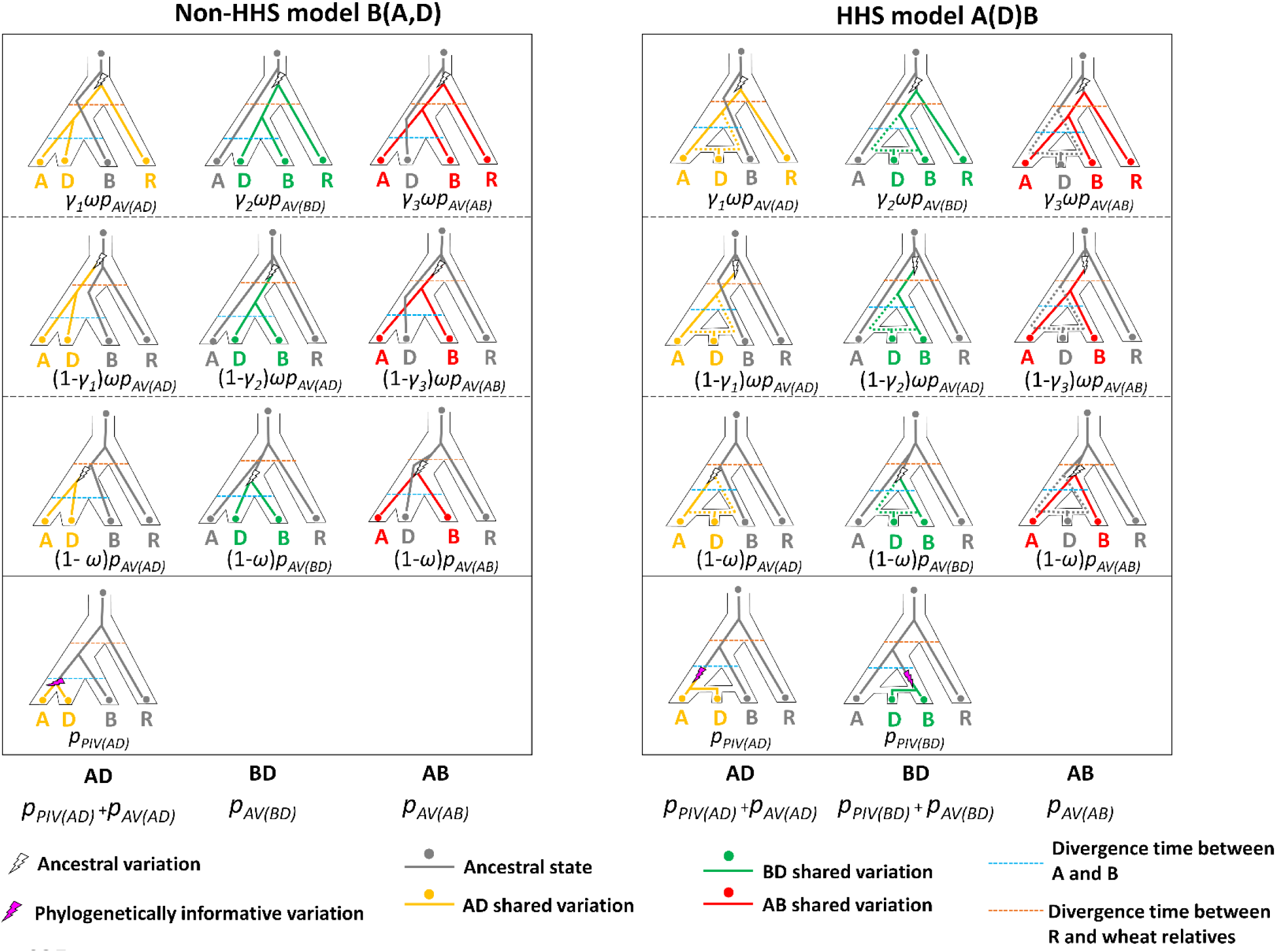
Difference in the probability of shared variations between the HHS and non-HHS models when rye is used as the outgroup. The probability of ancestral variations in shared variations is represented as *p*_*AV*_ and it is equal to *p*_*AV*(*AD*)_+*p*_*AV*(*BD*)_+*p*_*AV*(*AB*)_. Suppose ω is the proportion of AVs occurred prior to rye divergence, the probability of AVs before R (AVs^before R^) is *np*_*AV*_. The probability of AD, BD and AB types derived from AVs^before R^could be represented as ω*p*_*AV*(*AD*)_, ω*p*_*AV*(*BD*)_ and ω*p*_*AV*(*AB*)_, respectively. Suppose *γ*_1_, *γ*_2_ and *γ*_3_ are the proportions of shared variations between rye and those in AD, BD or AB from AVsbefore R, respectively, we have *p*_*ADR*_=*γ*_1_ω*p*_*AV*(*AD*)_, *p*_*BDR*_=*γ*_2_ω*p*_*AV*(*BD*)_ and *p*_*ABR*_ =*γ*_3_ω*p*_*AV*(*AB*)_. As the R genome has a similar genomic relatedness to the diploid A, B or D lineage (thus *γ*_1_, *γ*_2_ and *γ*_3_ are approximately equal), we thus have *p*_*ADR*_: *p*_*BDR*_: *p*_*ABR*_ = *p*_*AV*(*AD*)_: *p*_*AV*(*BD*)_: *p*_*AV*(*AB*)_.

**Supplementary Table 1.** The genomic relatedness between rye and each of the three subgenomes of bread wheat inferred from the shared SNPs in 1,614 orthologous genes^a^

| Data set<sup>b</sup> | Number of<br>SNPs | Shared SNPs |  |  | Genomic<br>relatedness<sup>c</sup> |
| --- | --- | --- | --- | --- | --- |
| A/B/R | 20530 | AB:70.5% | AR:14.7% | BR:14.8% | AB>>AR≈BR |
| A/D/R | 21044 | AD:74.5% | AR:12.9% | DR:12.5% | AD>>AR≈DR |
| B/D/R | 20566 | BD:73.8% | BR:13.3% | DR:12.9% | BD>>BR≈DR |
| A/B/D | 12170 | AB:26.7% | AD:38.9% | BD:34.4% | AD>BD>AB |

<sup>a</sup>The orthologous genes were identified from CDS of the three subgenomes of bread wheat, rye and barley (Supplementary data 4).
<sup>b</sup>The barley (Morex) genome was used as the outgroup to infer the ancestral states of a given variation;
<sup>c</sup>The results indicated that the rye genome shares a similar genetic relatedness with A, B or D subgenome of bread wheat.

**Supplementary Table 2.**
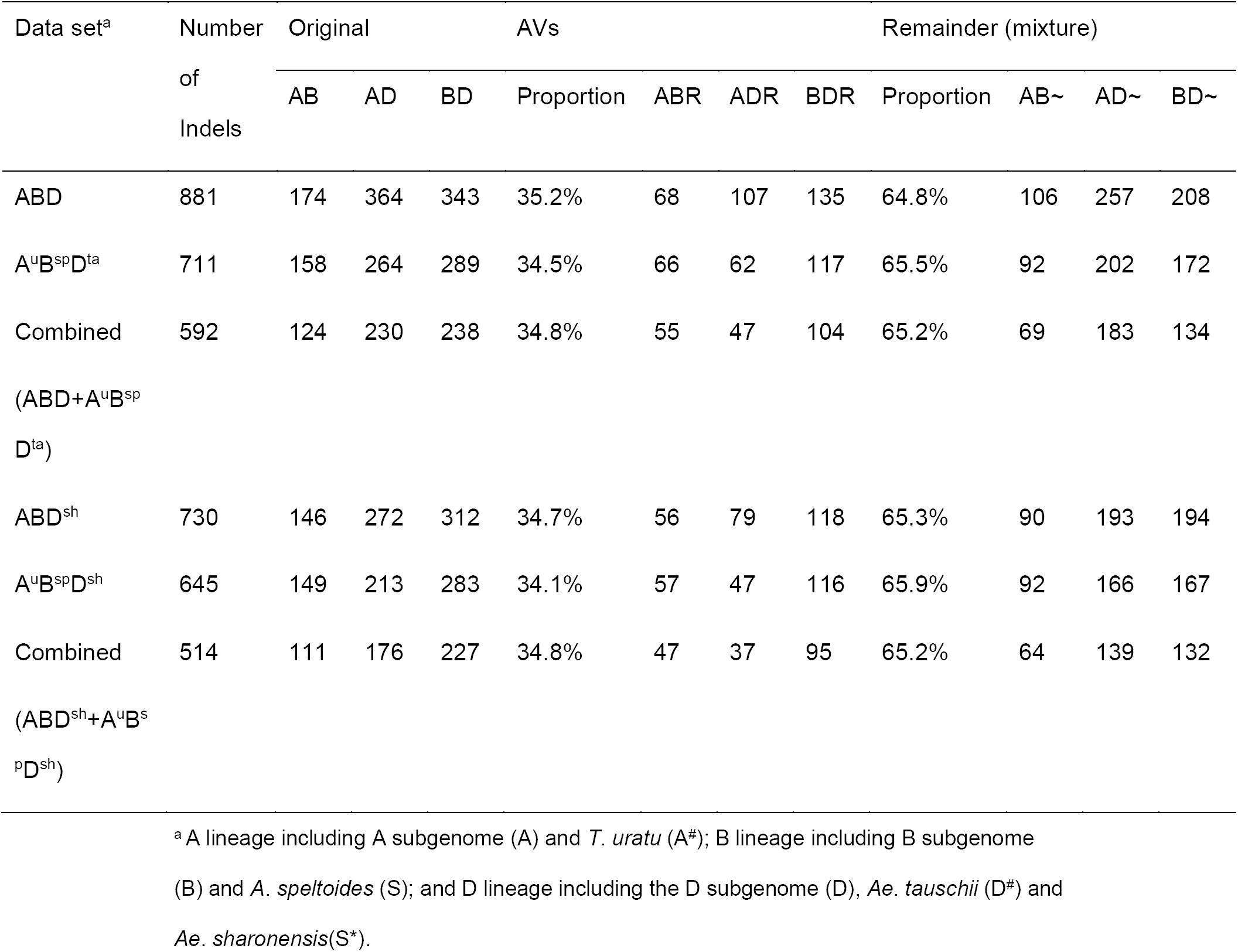
The distribution of shared Indels in the six different data sets

**Supplementary Table 3.**
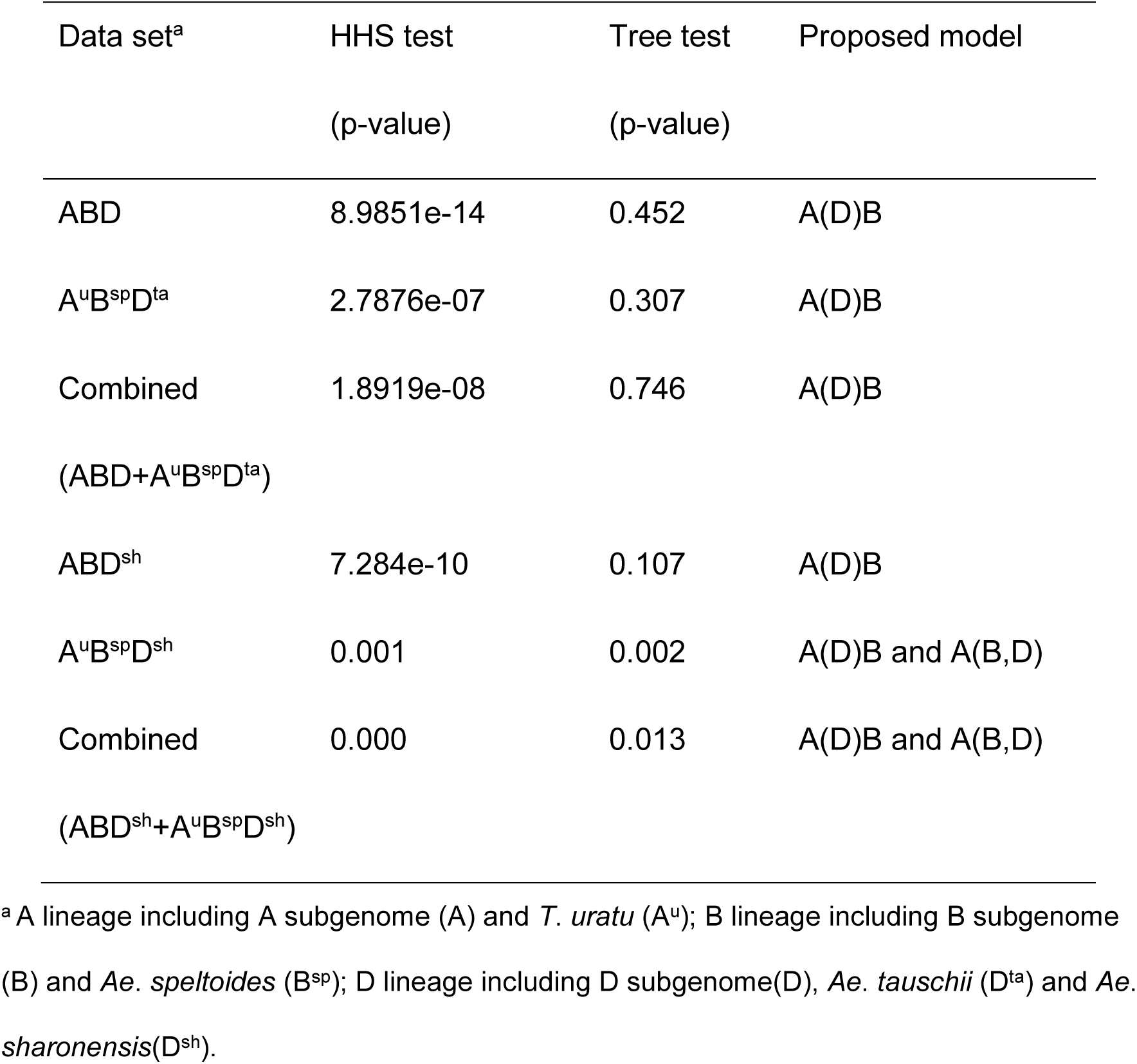
KKSC insertion significance test on the proposed phylogenetic models based on shared Indels

| Data set<sup>a</sup> | HHS test<br>(p-value) | Tree test<br>(p-value) | Proposed model |
| --- | --- | --- | --- |
| ABD | 8.9851e-14 | 0.452 | A(D)B |
| A<sup>u</sup>B<sup>sp</sup>D<sup>ta</sup> | 2.7876e-07 | 0.307 | A(D)B |
| Combined<br>(ABD+A<sup>u</sup>B<sup>sp</sup>D<sup>ta</sup>) | 1.8919e-08 | 0.746 | A(D)B |
| ABD<sup>sh</sup> | 7.284e-10 | 0.107 | A(D)B |
| A<sup>u</sup>B<sup>sp</sup>D<sup>sh</sup> | 0.001 | 0.002 | A(D)B and A(B,D) |
| Combined<br>(ABD<sup>sh</sup>+A<sup>u</sup>B<sup>sp</sup>D<sup>sh</sup>) | 0.000 | 0.013 | A(D)B and A(B,D) |

<sup>a</sup> A lineage including A subgenome (A) and *T. uratu* (A<sup>u</sup>); B lineage including B subgenome (B) and *Ae. speltoides* (B<sup>sp</sup>); D lineage including D subgenome(D), *Ae. tauschii* (D<sup>ta</sup>) and *Ae.* *sharonensis*(D<sup>sh</sup>).

**Supplementary Table 4.**
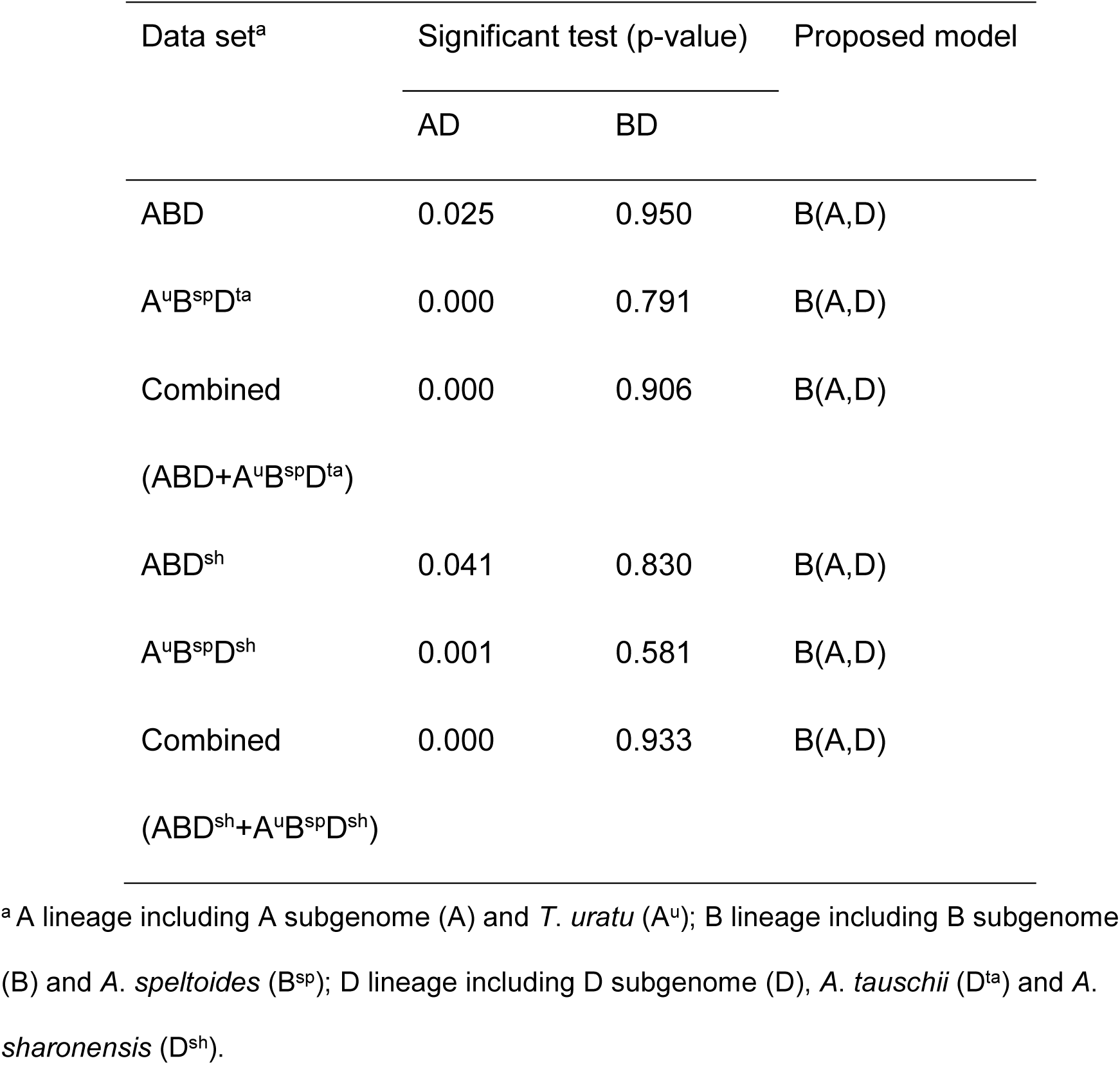
Significant test of PIV signals in AD and BD based on shared Indels

**Supplementary Table 5.**
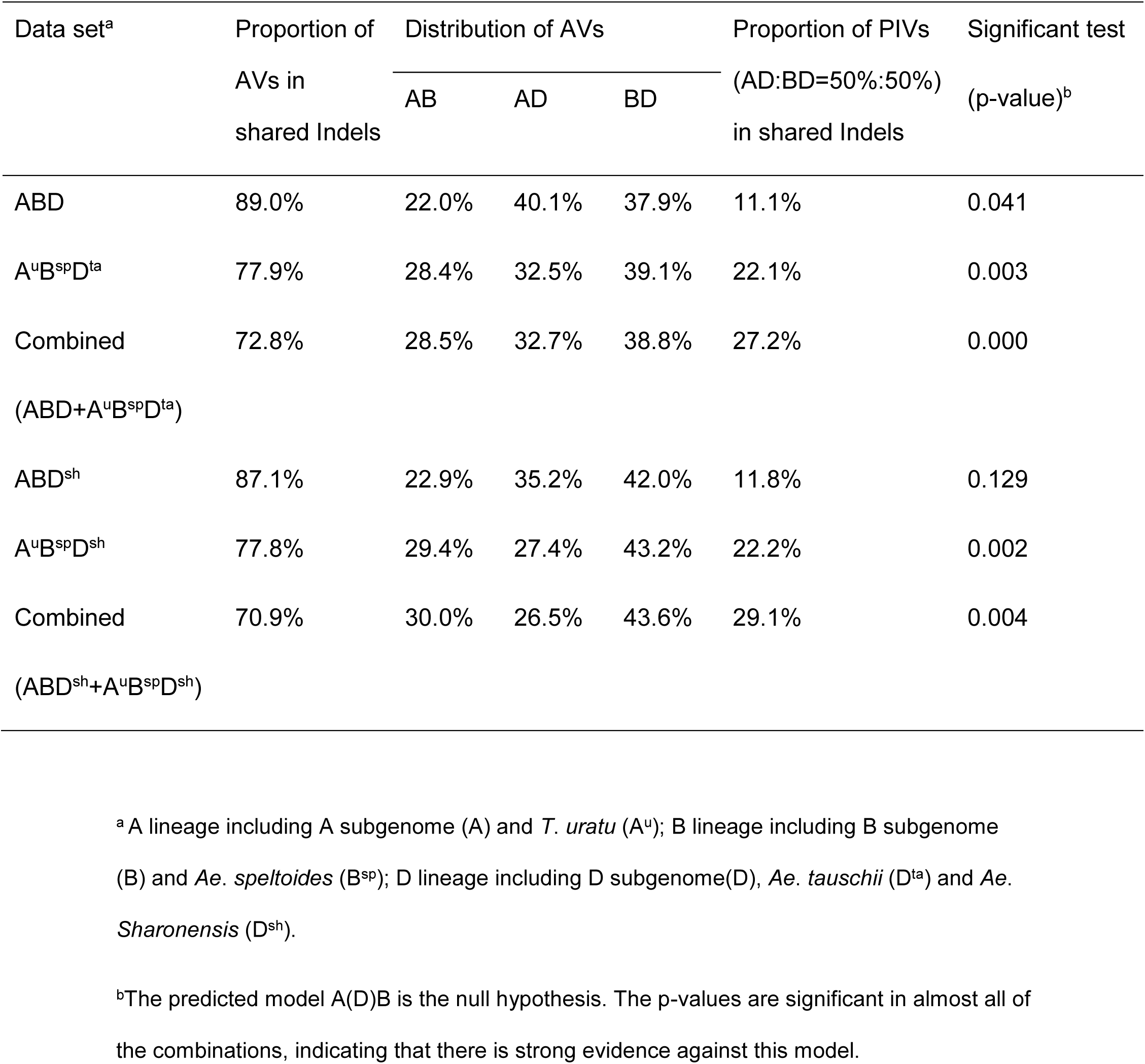
The statistical test for the HHS-model A(B)D (A:B=50%:50%).

| Data set<sup>a</sup> | Proportion of<br>AVs in<br>shared Indels | Distribution of AVs |  |  | Proportion of PIVs<br>(AD:BD=50%:50%)<br>in shared Indels | Significant test<br>(p-value)<sup>b</sup> |
| --- | --- | --- | --- | --- | --- | --- |
|  |  | AB | AD | BD |  |  |
| ABD | 89.0% | 22.0% | 40.1% | 37.9% | 11.1% | 0.041 |
| A<sup>u</sup>B<sup>sp</sup>D<sup>ta</sup> | 77.9% | 28.4% | 32.5% | 39.1% | 22.1% | 0.003 |
| Combined<br>(ABD+A<sup>u</sup>B<sup>sp</sup>D<sup>ta</sup>) | 72.8% | 28.5% | 32.7% | 38.8% | 27.2% | 0.000 |
| ABD<sup>sh</sup> | 87.1% | 22.9% | 35.2% | 42.0% | 11.8% | 0.129 |
| A<sup>u</sup>B<sup>sp</sup>D<sup>sh</sup> | 77.8% | 29.4% | 27.4% | 43.2% | 22.2% | 0.002 |
| Combined<br>(ABD<sup>sh</sup>+A<sup>u</sup>B<sup>sp</sup>D<sup>sh</sup>) | 70.9% | 30.0% | 26.5% | 43.6% | 29.1% | 0.004 |

<sup>a</sup> A lineage including A subgenome (A) and *T. uratu* (A<sup>u</sup>); B lineage including B subgenome (B) and *Ae. speltoides* (B<sup>sp</sup>); D lineage including D subgenome(D), *Ae. tauschii* (D<sup>ta</sup>) and *Ae.* *Sharonensis* (D<sup>sh</sup>).
<sup>b</sup>The predicted model A(D)B is the null hypothesis. The p-values are significant in almost all of the combinations, indicating that there is strong evidence against this model.

